# GFP transformation sheds more light on a widespread mycoparasitic interaction

**DOI:** 10.1101/516781

**Authors:** Márk Z. Németh, Alexandra Pintye, Áron N. Horváth, Pál Vági, Gábor M. Kovács, Markus Gorfer, Levente Kiss

**Affiliations:** Plant Protection Institute, Centre for Agricultural Research, Hungarian Academy of Sciences (MTA-ATK), Budapest, H-1525, Hungary; Eötvös Loránd University, Institute of Biology, Department of Plant Anatomy, Budapest, H-1117, Hungary; AIT Austrian Institute of Technology; Konrad Lorenz Straße 24, A-3430 Tulln, Austria; University of Southern Queensland, Institute for Life Sciences and the Environment, Centre for Crop Health, Toowoomba, Qld 4350, Australia

## Abstract

Powdery mildews (PMs), ubiquitous obligate biotrophic plant pathogens, are often attacked in the field by mycoparasitic fungi belonging to the genus *Ampelomyces*. Some *Ampelomyces* strains are commercialized biocontrol agents of crop pathogenic PMs. Using *Agrobacterium tumefaciens-mediated* transformation (ATMT), we produced stable *Ampelomyces* transformants that constitutively expressed the green fluorescent protein (GFP), to (i) improve the visualization of the PM-*Ampelomyces* interaction; and (ii) decipher the environmental fate of *Ampelomyces* before and after acting as a mycoparasite. Detection of *Ampelomyces* structures, and especially hyphae, was greatly enhanced when diverse PM, leaf and soil samples containing GFP transformants were examined with fluorescence microscopy compared to brightfield and DIC optics. We showed for the first time that *Ampelomyces* can persist up to 21 days on PM-free host plant surfaces, where it can attack PM structures as soon as these appear after this period. As a saprobe in decomposing, PM-infected leaves on the ground, and also in autoclaved soil, *Ampelomyces* developed new hyphae, but did not sporulate. These results indicate that *Ampelomyces* occupies a niche in the phyllosphere where it acts primarily as a mycoparasite of PMs. Our work has established a framework for a molecular genetic toolbox for *Ampelomyces* using ATMT.

## 1. Introduction

Mycoparasites, i.e. fungi that parasitize other fungi, are commonly found in most terrestrial ecosystems, the best known species being those that attack fungal plant pathogens^1–4^. A number of mycoparasites have been long studied and commercially utilized as biocontrol agents (BCAs) of crop pathogens^3,4^; others have been in focus as components of natural multitrophic relationships^2,5–9^. Direct observation of interfungal parasitic relationships is notoriously difficult at cellular or hyphal level, using classical light, scanning and transmission electron microscopy (LM, SEM and TEM) and other visualization methods^1,10–13^. Recombinant DNA technologies, such as those leading to the production of transformed strains that constitutively or inducibly express fluorescent proteins, e.g., the green fluorescent protein (GFP), have been applied in the study of mycoparasitism since these technologies became available in fungal biology research^14^. For example, experiments using GFP-expressing BCAs, such as *Clonostachys* and *Trichoderma* strains, resulted in improved visualization of mycoparasitic interactions between these fungi and their plant pathogenic mycohosts^15–17^. Bitsadze et al.^18^ visualized the degradation of *Sclerotinia sclerotiorum* sclerotia separately, and also simultaneously, by two transformed BCAs, *Microsphaeropsis ochracea* and *Paraphaeosphaeria minitans*, expressing red fluorescent protein (DsRed) and GFP, respectively. The attack of the cultivated mushroom *Agaricus bisporus* by *Lecanicillium fungicola*, has also been studied with GFP-expressing mycoparasitic strains^19^.

A remarkable example of a widespread natural interfungal parasitic relationship is the interactions between powdery mildews (PMs), common obligate biotrophic plant pathogens of more than 10,000 angiosperms, including important crops^20,21^, and pycnidial fungi belonging to the genus *Ampelomyces*^22^. This intricate interaction has not been examined previously by fluorescent protein biotechnology. In fact, this is the oldest known interfungal parasitic relationship, first revealed by De Bary in the 19^th^ century using simple LM^22–24^, and later confirmed by detailed LM, SEM and TEM studies^25–28^. As De Bary^23^ and subsequent studies have shown, one-celled conidia of *Ampelomyces* are released from pycnidia produced inside the PM mycelium by the rupture of the pycnidial wall; conidia then germinate on the host plant surfaces, penetrate PM hyphae found in their vicinity and invade these internally, growing from cell to cell through the septal pores of the mycohost^25,26^. The early stage of mycoparasitism is biotrophic^28^, but the invaded hyphal cells later begin to die, and a necrotrophic interaction results^26^. This process ends with intracellular sporulation of *Ampelomyces*, i.e. production of pycnidia mostly inside PM conidiophores, immature conidia, and, if present, also in immature chasmothecia^25–31^ (sexual fruiting bodies of PMs).

Parasitized PM colonies continue their growth on host plant surfaces, but their asexual and sexual sporulation is reduced, or completely stopped, by Ampelomyces^29–34^. A transcriptomic study revealed the upregulation of some genes related to toxin biosynthesis, together with other potentially mycoparasitism-related proteins, such as secreted proteases and putative virulence factors during mycoparasitism^35^. However, approx. 50% of the *Ampelomyces* transcripts did not relate to any known protein sequences^35^. This may indicate that a part of the *Ampelomyces* proteome is unique, or this type of mycoparasitism has not been studied in sufficient detail in other interfungal parasitic relationships.

The natural occurrence of *Ampelomyces* has been globally reported in dozens of PM species^9,29,36^. Phylogenetic analyses have revealed that *Ampelomyces* strains isolated from diverse PM species, and cultured on agar media, belong to distinct lineages that are not clearly associated with their original mycohosts^9,37–42^. All *Ampelomyces* strains tested were able to parasitise several PM species, in addition to their original mycohosts, both in the laboratory^28,31,33,34,37,38^ and in field experiments^31,32,40^, after being subcultured on agar media. A formal taxonomic revision of the genus is still warranted^31^. PM species are each specialized to one or a few host plant species^20,21^, while genetically distinct *Ampelomyces* strains seem to be their generalist mycoparasites^28,31,40,41,43^. These interactions are interesting natural tritrophic relationships^2^, and have been included in a number of ecological studies^5–8^. As PMs are themselves parasites, *Ampelomyces* strains are also called as hyperparasites^2,6–8,22^. A number of *Ampelomyces* strains have been exploited and commercialized as BCAs of PMs infecting grapes and vegetables^24,29–32^.

In the absence of mycohosts, i.e. living PM colonies, the activity of *Ampelomyces* strains in the field is an interesting and debated question. Brimner & Boland^44^ suggested that the application of *Ampelomyces* as a BCA may result in unwanted interactions with soil fungi, as non-target effects of biocontrol procedures, although there is no evidence for such interactions^45^. It has been repeatedly shown that several strains identified as *Ampelomyces* were confused with *Phoma-like* pycnidial fungi isolated from diverse environments^37,46,47^, which has led to misconceptions about the biology of *Ampelomyces*^22,24,47^. The only experimental evidence for survival of *Ampelomyces* in the environment, in the absence of living PM colonies, was the detection of a number of overwintering strategies linked to PM structures in temperate regions^29,40,48^. As *Ampelomyces* strains can be cultured on artificial media^9,29,30,32,39,41,42^, the possibility that these fungi persist in the environment as saprobes cannot be excluded. It is not known whether these mycoparasites develop further as saprobes in senescent leaves that were once infected with *Ampelomyces*-parasitized PMs, as hypothesized by De Bary^23^ and Yarwood^49^ several decades ago, or whether *Ampelomyces* interacts with soil-borne fungi after those leaves have been decomposed at the soil level.

It is also not known what happens to conidia of *Ampelomyces* on PM-free aerial plant surfaces after being naturally released and splash-dispersed from pycnidia produced in already parasitized PM colonies, or sprayed on crops as BCAs. Clearly, a method to better visualize *Ampelomyces* structures inside and outside PM colonies, such as the use of GFP-expressing mutants in laboratory experiments, would greatly enhance the study of environmental fate of *Ampelomyces* in the absence of their mycohosts, i.e. before, after, or instead of acting as a mycoparasite, and would also shed more light on their development inside PM colonies. The objectives of this work were to (i) transform *Ampelomyces* strains to produce mutants that constitutively express GFP in culture; (ii) characterize transformants in terms of transgene copy numbers; (iii) characterize transformants in terms of morphology and growth in culture as well as mycoparasitic activity compared to the wild strains; (iv) test whether GFP expression of selected transformants is stable during mycoparasitic development inside PM structures; (v) compare the visualization of the mycoparasitic interaction with GFP expression, using fluorescence microscopy, versus other optical microscopy techniques; (vi) determine whether *Ampelomyces* conidia can survive for at least 2-3 weeks on PM-free host plant surfaces; (vii) observe the fate of *Ampelomyces* structures during and after senescence of leaves infected with *Ampelomyces*-parasitized PM; and (viii) examine if there is any saprotrophic development of *Ampelomyces* in autoclaved soil.

## 2. Materials and Methods

### 2.1. Fungal and plant materials

Recently, a comprehensive screening compared the mycoparasitic activity of 33 *Ampelomyces* strains in grape PM (*E. necator*) as well as their growth and sporulation rate in culture^31^. Two among those 33 strains, RS1-a and GYER, were used in this study. RS1-a was originally isolated from *P. pannosa* infecting rose (*Rosa* sp.) and GYER from *E. arcuata* infecting hornbeam (*Carpinus betulus*). RS1-a was included in this study because it was the best performing strain in terms of mycoparasitic activity in grape PM and sporulation in culture^31^, while GYER was selected for this work to test the transformation protocol on another distinct *Ampelomyces* genotype, as well. The two stains were maintained on Czapek-Dox medium supplemented with 2% malt (MCzA) as described earlier^31^.

Mycoparasitic tests were performed with the following five PM species, maintained in the greenhouse on their respective host plants, grown in pots: *E. necator* infecting grapevine (*Vitis vinifera* cv. Chardonnay), *P. xanthii* infecting cucumber (*Cucumis sativus* cv. Rajnai Fürtös); *B. graminis* f. sp. *hordei* infecting barley (*Hordeum vulgare* MW08-16), *P. neolycopersici* infecting tomato (*Solanum lycopersicum* cv. Kecskeméti Jubileum), and *L. taurica* infecting pepper (*Capsicum annuum* cv. Total). Potted grapevine plants continuously produced young leaves that supported the maintenance of *E. necator* in the greenhouse throughout the season. Potted cucumber, barley, tomato and pepper plants, infected with their respective PMs, had to be replaced with newly grown and freshly inoculated plants every 3 to 6 weeks to ensure the continuous maintenance of the other four PM species in the greenhouse.

### 2.2. Phylogenetic analyses

An approx. 850 bp long part of the actin gene (*act1)* of GYER was amplified and sequenced as described previously^41^, and deposited in GenBank under accession number MH879022. This sequence was aligned with the *act1* sequence dataset used by Pintye *et al*.^41^ using MAFFT online^56^ with FFT-NS-i algorithm. The rDNA internal transcribed spacer (ITS) sequence dataset used by Pintye *et al*.^41^ was supplemented with the ITS sequence of GYER, determined earlier^40^, and was aligned with MAFFT online using E-INS-i. These alignments were combined using MEGA6^57^ to produce a concatenated ITS_act1 dataset. The final alignment (TreeBase study ID 23290) was 1333 characters long, consisting of 502 characters for the ITS and 831 for the *act1* dataset. This combined alignment was introduced to RaxML raxmlGUI 1.5^58,59^ and a maximum likelihood analysis was conducted using GTR+G substitution model and ML estimation of base frequencies. Partitions were set to correspond to the two loci. Supports of the branches were calculated from 1,000 bootstrap replicates. The tree with the highest likelihood value was visualised in MEGA6^57^ and TreeGraph 2.13.0^60^. *Phoma herbarum* CBS567.63 was used as outgroup based on a previous work^41^.

### 2.3. Transformation of *Ampelomyces* strains

As hygromycin B (Sigma-Aldrich) was used as the selection marker during transformations, first, as a preliminary step, we determined the lowest concentration that causes complete growth inhibition of strains RS1-a and GYER in culture. The following concentrations of hygromycin B added to malt extract agar (MEA; Merck) were tested: 5 mg/l, 10 mg/l, 25 mg/l, 50 mg/l and 100 mg/l. Each concentration was tested by inoculating three plates with a 5 mm diameter coeval mycelial disc of one of the two strains. These tests, done twice, revealed that 50 mg/l hygromycin B completely inhibited the growth of both strains. Therefore, this concentration was used in the subsequent work.

Transformation of the two strains was done based on previously published protocols^50,61^ as follows. First, 15 to 20 small fungal colony fragments, approx. 2-4 mm x 2-4 mm, were transferred to sterile cellophane sheets placed on MEA in 9 cm diameter plates. Five such plates were prepared for RS1-a and another ten for GYER. Strains were grown for four days in dark at 23 °C, then the cellophane sheets bearing the growing fungal colonies were transferred to plates containing Moser induction medium^50^ (MoserIND).

*Rhizobium radiobacter* (previously known as *Agrobacterium tumefaciens*) strain AGL1^62^ carrying the binary vector pCBCT^50^ was used in the transformation work. This vector contains the *SGFP* gene with *toxA* promoter^14^ and the *hph* gene driven by *trpC* promoter^63^. Both promoters provide a stable constitutive expression of the respective genes^14^.

Bacteria were grown overnight at 28°C in 30 ml pastic tubes under continuous agitation (180 rpm) in a shaker in 4 ml tryptic soy broth supplemented with 50 μg/ml kanamycin. Bacteria were then pelleted by centrifugation for nine min at 3,800 g and resuspended in *Agrobacterium* Induction Medium^50^ (AtIND). Their induction was reached during growth for 6 h under continuous agitation at 180 rpm at 28°C. Aliquots of the induced bacterial culture were pipetted directly on *Ampelomyces* colonies growing on cellophane sheets placed on MoserIND. Co-culture plates were incubated for four days at room temperature, then cellophane sheets from these plates were transferred to 9 cm diameter plates containing 25-28 ml selective medium each (MEA with 50 mg/l hygromycin B and 100 mg/l cefotaxime, Duchefa Biochemie) and incubated in the dark at 22°C for 4 to 8 weeks, until emergence of visible fungal colonies. Hyphae and, if produced, conidia of actively growing putative transformants were illuminated with blue light and colonies exhibiting green fluorescence were transferred each to a new 6 cm diameter plate with selective medium. Colonies exhibiting fluorescence when excited with blue light were considered as GFP transformants of the respective *Ampelomyces* strains. To produce monoconidial transformants, 1 ml sterile water with 50 mg chloramphenicol was pipetted onto each plate containing a sporulating colony of a RS1-a transformant and the colony surface was rubbed with a sterile artist’s brush to release as many conidia as possible from pycnidia. Conidial suspensions, diluted 10^4^-10^6^ times with sterile water, were spread on new 6 cm diameter plates containing MCzA. A similar procedure was carried out with the actively growing colonies of GYER transformants; suspensions produced from non-sporulating GYER colonies were diluted 100x, and hyphal fragments of these suspensions became the colony forming units. Plates were incubated for one week at room temperature, then the emerged individual colonies were transferred each to a new plate with MCzA, and three weeks later subcultured on the same medium. GFP expression of hyphae and, if produced, conidia of the subcultured colonies was verified with fluorescence microscope. These all exhibited green fluorescence under blue light (Fig. 1), thus were considered as GFP transformants of the wild type strains RS1-a and GYER, respectively. Seven transformants of RS1-a and six transformants of GYER were used in subsequent work; their designations are shown in Supplementary Table S1. These 13 transformants were subcultured every 4-6 weeks in 6 cm diameter plates on MCzA. GFP expression of hyphae was verified with fluorescent microscopy before every subculturing. Hygromycin B was not added to MCzA used to maintain the transformants in culture as these remained stable without selection pressure during the duration of this work.

**Figure 1.**
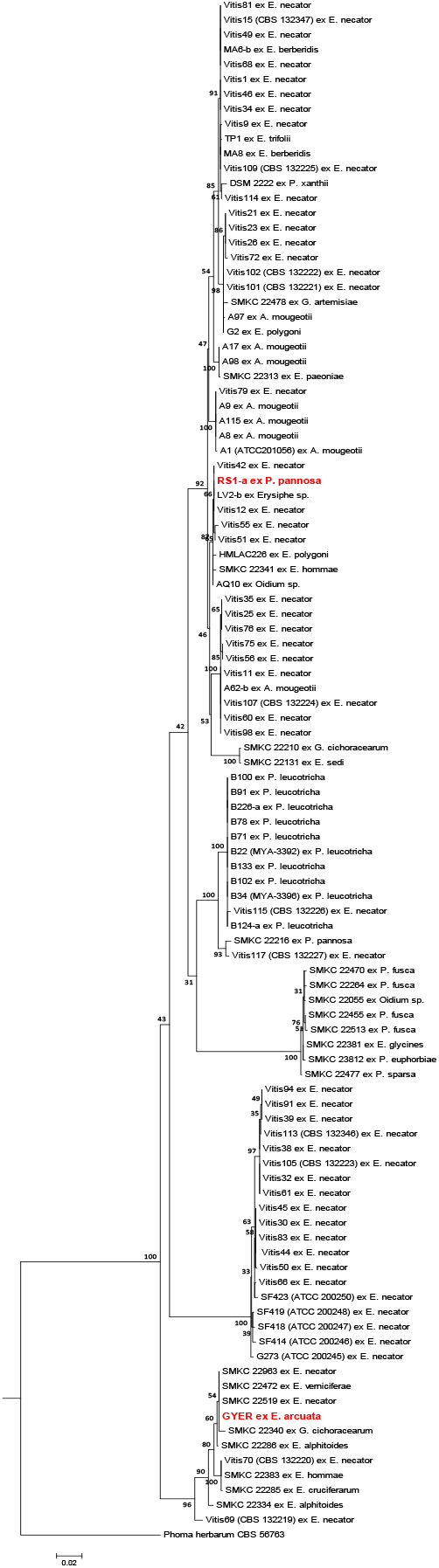
Phylogenetic tree with the highest likelihood value based on a concatenated dataset consisting of nrDNA ITS and partial actin gene (*act1*) sequences of 103 *Ampelomyces* strains, including RS1-a and GYER (both in red), the two strains used in the transformation work reported here. A concatenated ITS and *act1* sequence of *Phoma herbarum* strain CBS567.63 was used as outgroup. Numbers above the branches show bootstrap support values calculated from 1,000 replications. Support values below 70% and in subclades are not shown. Bar indicates 0.02 expected change per site per branch. Designations, place and date of collection, and GenBank accession numbers of both ITS and *act1* sequences of the *Ampelomyces* strains that were included in the analysis, but not in the transformation work, were published in earlier studies^39,41^.

### 2.4. PCR-verification of T-DNA insertion into transformants

DNA was extracted from the 13 transformants as described earlier^64^, with centrifugation lengths extended to 20 min. Insertion of transfer DNA (T-DNA) in the transformants’ genomes was checked by amplification of two fragments of T-DNA, *hph* and *SGFP*, using primers hph-F1/hph-qR2 and GFPF/GFPR^17^. Primers hph-F1 and hph-qR2 were designed based on the pCBCT sequence^50^ using SnapGene Viewer software (version 4; GSL Biotech). In these amplifications Phusion High-Fidelity DNA Polymerase (Thermo Fischer Scientific) was used according to the manufacturer’s recommendations. The PCR program was as follows: 98°C denaturation for 2 minutes, followed by 35 cycles each consisting of 98°C for 10 sec, 57°C (for hph) or 66°C (for *SGFP*) for 10 sec, and 72°C for 30 sec, and concluded with a final extension at 72°C for 5 minutes. Negative controls (ultra-pure water) and non-template controls (DNA from the respective wild type strains) were included in all amplifications. The resulting PCR products were separated on 1% agarose gel and visualized using GelRed Nucleic Acid Gel Stain (Biotium) under UV illumination in GelDoc-It system (UVP). Amplicons were sent for sequencing to LGC Genomics GmbH (Germany). Sequencing was done with primers hph-qF1 and GFPF, respectively. Resulting sequences were aligned using MEGA7^65^ using default settings. Primer sequences are shown in Supplementary Table S3.

### 2.5. Determination of T-DNA insert copy numbers

Copy number of T-DNA in transformants was determined by quantitative real time PCR (qPCR) using the comparative Ct (2^-ΔΔCt^) method^66^, the most robust method for this purpose^67^. This approach has already been shown to work well in fungal transformants^68^ and it is based on the relative quantification of copy numbers of an unknown target region compared to a known copy number internal reference gene. We targeted *hph* of T-DNA as the unknown copy number region. A putatively single copy gene, *euknr*, the eukaryotic nitrate reductase encoding gene^69^ was sequenced in wild type RS1-a and GYER strains with the Ascomycetes-specific nested PCR method described previously^69^. The newly obtained sequences were deposited in GenBank under accession numbers MH879020 and MH879021. A BLAST search with these sequences on the publicly available *Ampelomyces* genome (https://genome.jgi.doe.gov/Ampqui1/Ampqui1.home.html) using the default settings of the webpage resulted in a single hit, indicating that *euknr* is a single copy gene in *Ampelomyces*. Thus, *euknr* was used as a single copy internal reference gene in qPCR measurements.

To produce a calibrator sample for the comparative Ct method, the *hph* gene of pCBCT and an approximately 1 kb fragment of *euknr* of RS1-a and GYER were amplified with Phusion Green Hot Start II High-Fidelity PCR Master Mix (Thermo Fisher Scientific). Primers trpCP02/CB02^50^ were used to amplify *hph* and niaD31F (for RS1-a) or niaD31F-alt (for GYER)/niaD32R to amplify *euknr*. Primers niaD31F, niaD31F-alt and niaD32R were designed with SnapGene Viewer software using the *euknr* sequences determined earlier to allow direct amplification of *euknr* from our strains. PCR products were purified with QIAquick PCR Purification Kit (Qiagen) and the fragment containing *hph* was subjected to a treatment with Fast DNA End Repair Kit (Thermo Fisher Scientific) to produce 5’ phosphorylated DNA for blunt end ligation. Resulting treated DNA was purified with QIAquick PCR Purification Kit. Ligation was done with a Rapid DNA Ligation Kit (Thermo Fisher Scientific) and the resulting ligation mixture was used as a target for PCR with primers trpCP02/niaD32R to amplify the targeted *hph_euknr* fusion product. The product was purified from 1% agarose gel with GenElute Gel Extraction Kit (Sigma-Aldrich) and cloned into pJET1.2/blunt vector using CloneJET PCR Cloning Kit (Thermo Fisher Scientific). Resulting plasmid products were transformed into One Shot TOP10 Chemically Competent *E. coli* cells (Invitrogen). One positive clone was selected for *hph* fused to *euknr* of RS1-a, and another one for *hph* fused to *euknr* of GYER. These clones were grown separately overnight in lysogeny broth (LB) medium and used for plasmid extraction with GenElute Plasmid Miniprep Kit (Sigma-Aldrich). PCR amplifications, product purifications, phosphorylation, ligation, cloning and plasmid extraction were conducted according to the manufacturers’ protocols. Plasmid solutions of approx. 5 x 10^5^-10^6^ copies/μl were produced, and used in qPCR as 1:1 copy number calibrators.

For qPCR measurements DNA was extracted from the 13 transformants using DNeasy Plant Mini Kit (Qiagen). qPCR primers were designed manually with the aid of SnapGene Viewer software. Primers hph-qF2 and hph-qR2 were designed based on the pCBCT sequence^50^ while primers qNR-F2/qNR-R2 were designed based on the *euknr* sequences of RS1-a and GYER determined in this study. First, these primer sets were tested during qPCRs using dilution series (undiluted DNA, and 2x, 5x, 10x, 50x and 100x dilutions in ultra-pure water) of RS1-a transformant A2 to determine efficiency and range of linearity. Reactions were run in triplicate in a Bio-Rad CFX96 Touch C1000 qPCR machine in 10 μl final volumes with 5 μl iTaq Universal SYBR Green Supermix (Bio-Rad), 0.5 μl of each primer and 1 μl DNA extract, using the following PCR program: denaturation at 95°C for 5 min, followed by 30 cycles each consisting of 95°C for 10 sec, 59°C for 10 sec, and 72°C for 30 sec. Specificity of the reactions was checked by registering a melt curve. A negative control reaction has always been included in each qPCR. Measured qPCR efficiencies were ~90% and ~89%, respectively and all the tested dilutions were found to fit the linear range. Correlation coefficients (R^2^) of regression curves were 0.992 for *hph* and 0.998 for *euknr* amplifications. The same qPCR protocol and reaction composition were used for copy number measurements using plasmid extracts and transformants’ DNA samples as targets. Copy numbers were calculated using the comparative Ct (2^-ΔΔCt^) method^66^. Primer sequences are shown in Supplementary Table S3.

### 2.6. Morphology and saprobic growth characteristics of transformants

The morphology of the coeval colonies of the seven RS1-a and the six GYER transformants, grown on MCzA, as well as the morphology of their pycnidia and conidia produced in culture, and examined with DIC optics, were compared to those of the wild type RS1-a and GYER strains, to observe if any obvious morphological differences were developed as a result of the transformation. Saprotrophic growth of transformants was measured during growth tests at 22°C as described previously^31^ with eight replicates of each transformant. High resolution photos of four-week old colonies were used to measure the area of each fungal colony using ImageJ software version 1.51^70^.

### 2.7. Setup of mycoparasitic tests with transformants

As the wild type GYER strain and its transformants exhibited unreliable sporulation patterns during this study, these fungi were excluded from PM inoculation experiments. Thus, only the wild type RS1-a strain and its seven transformants were used to test if mycoparasitic activity was altered in terms of penetration of PM hyphae, intrahyphal growth and production of intracellular pycnidia, as a result of transformation. Young, sporulating *P. xanthii* colonies on potted cucumber plants were inoculated with conidial suspensions, 10^6^ conidia/ml, prepared from each of the seven RS1-a transformants and the wild type as described earlier^37^. Conidial suspensions were sprayed each on three plants until runoff. To verify the viability/GFP expression of *Ampelomyces* conidia used in these treatments, 30 μl of each conidial suspension was spread immediately after inoculations in a 6 cm diameter plate containing 1.5% water agar medium covered by one layer of sterile cellophane. Plates were incubated at room temperature for 24 h, then cellophane pieces bearing fungal material were cut out, placed on a slide, and examined with a fluorescence microscope as detailed below. Inoculated plants were kept in closed transparent isolation chambers for ten days at 22°C, 80-90% relative humidity (RH), and 16 h daily illumination. Three PM-infected plants sprayed with water until runoff, and kept in another isolation chamber, served as negative controls. Samples of the PM mycelia treated with *Ampelomyces*, or water in case of negative controls, were microscopically examined as described below. The experiment was carried out three times: four, five and then 30 months after production and repeated subculturing of the transformants.

### 2.8. Microscopy

Parts of the PM mycelia treated with *Ampelomyces* were removed from leaves with 3-5 cm long pieces of crystal clear cellotape. These were placed on microscope slides, in droplets of distilled water or 10% glycerol, without being covered with coverslips, and first examined with brightfield, DIC and/or phase contrast microscopy with a Zeiss Axioskop 2 Plus microscope. GFP fluorescence was detected in epifluorescent mode using a fluorescent filter set composed of 450-490 nm excitation filter, 495 nm dichroic mirror and 500-550 nm barrier filter. Photographs were taken with Zeiss AxioCam ICc5 camera using Zeiss ZEN 2011 software (Carl Zeiss Microscopy GmbH). Adobe Photoshop was used for minor colour and contrast corrections on microphotographs, without any content changes.

To prepare samples for CLSM, one or more intracellular pycnidia of *Ampelomyces* together with a small part of the parasitized PM mycelium were removed with glass needles from leaves under a dissecting microscope, and placed on a microscope slide, in a droplet of water, to be first examined with brightfield microscopy. CLSM observations were done using a Carl Zeiss 410 LSM as described previously^71^. Two μm thick optical slices were recorded and 10-20 slices were used to compile projections.

### 2.9. Mycoparasitic activity of RS1-a transformants in *P. xanthii* colonies on cucumber

To compare mycoparasitic activities, i.e. penetration of PM hyphae, intrahyphal growth and intracellular pycnidial development, of the seven RS1-a transformants to the wild type strain in the experimental setup described above, PM colonies were sampled with cellotape 10 days after inoculations with *Ampelomyces*, and examined with brightfield, DIC and fluorescence microscopy, together with samples taken from PM-infected plants that served as negative controls. CLSM observations were also performed. At the same time, to quantify the levels of mycoparasitism, nine individual PM colonies were randomly selected on three cucumber leaves for each transformant and the wild type strain, and the number of intracellular pycnidia produced in conidiophores of *P. xanthii* was determined under a stereomicroscope in each colony, in a 1.25 mm^2^ area, using a method developed earlier^40^.

### 2.10. Statistical analyses

Statistical analyses were conducted using IBM SPSS 16.0. All the results were expressed as the mean ± standard deviation (s.d.) data of eight replicates. Data determined during colony growth tests in culture and quantification of mycoparasitic activities of transformants and the wild type strain in *P. xanthii* on cucumber were checked for normality of the data and homogeneity of variances using Kolmogorov-Smirnov test and Levene’s test, respectively. The transformants’ values were compared to those of the wild type used as control in one-way ANOVA (with an alpha value of 0.05) coupled with either two-sided Dunnett or Dunett T3 post hoc test. All the analyses were run with 95% confidence intervals. Descriptive statistics and additional details of statistical analyses are given in Supplementary Table S4.

### 2.11. Mycoparasitic activities of transformants in other PM fungi

To check the effectiveness of GFP expression of RS1-a transformants in the visualisation of mycoparasitic activity of *Ampelomyces* in PMs other than *P. xanthii* infecting cucumber, mycoparasitic tests were carried out with *E. necator* infecting grapevine, *B. graminis* infecting barley, *P. neolycopersici* infecting tomato and *L. taurica* infecting pepper according to the setup described above. Transformants used in these tests are listed in Supplementary Table S2. Each experiment was carried out at least twice. Brightfield, phase contrast and DIC microscopy as well as CLSM were used to examine mycelial samples taken at the end of each experiment. One of the transformants, B3, was re-isolated from *P. xanthii* colonies as described earlier^38^, and checked for GFP fluorescence and hygromycin B resistance on selective medium.

### 2.12. Testing survival and potential development of *Ampelomyces* on cucumber in the absence of the mycohost

To examine whether *Ampelomyces* can survive for up to three weeks on PM-free host plant surfaces, conidial suspensions of the wild type RS1-a strain and those of two transformants, C10 and G4, were prepared, and the viability/GFP expression of *Ampelomyces* conidia were checked on 1.5% water agar medium as described above. Conidial suspensions from these three *Ampelomyces* materials were each sprayed until runoff on (i) 42 potted healthy cucumber seedlings with three true leaves each, produced in a PM-free growth chamber; these plants were infected with PM sequentially, four to 21 days after inoculation, in groups of six, as detailed below; (ii) six other healthy cucumber seedlings from the same plant cohort, to monitor the development of *Ampelomyces* in the absence of PM for up to 21 days; and (iii) six older cucumber plants, with six to eight true leaves each, heavily infected with PM, to serve as positive controls. Six healthy seedlings from the cohort with three true leaves served as negative controls, being sprayed with water. Each plant was kept in isolation, under a transparent cover, at 25°C, 80-90% RH and 16 h daily illumination until the end of the experiment that was carried out twice.

To examine whether *Ampelomyces* can parasitize PM colonies after being present on PM-free plant surfaces for up to three weeks, four, seven, 10, 12, 14, 18 and 21 days after treatments with RS1-a, C10, G4 and water, respectively, six plants per treatment were infected with *P. xanthii* by dusting conidia onto their leaves from coeval PM colonies maintained on other potted plants in the greenhouse. The young PM mycelium developed on each plant was examined with brightfield, DIC and fluorescence microscopy 10 days after infection for the presence of *Ampelomyces* structures, including GFP expressing pycnidia on leaves treated with C10 and G4.

To monitor the fate of *Ampelomyces* on PM-free cucumber seedlings, approx. 2 x 2 cm areas of the inoculated leaf surfaces, two per plant and treatment, were sampled ten, 14 and 21 days following treatments, with cellotape pieces, gently applied, and then removed from the treated leaves with a forceps. Cellotape pieces were examined with brightfield and fluorescence microscopy to detect germination and further development of *Ampelomyces* conidia.

The presence of intracellular pycnidia of RS1-a, C10 and G4 on the PM-infected plants used as positive controls was also examined with brightfield and fluorescence microscopy.

### 2.13. Testing survival and potential development of *Ampelomyces* in senescent/decomposed PM-infected leaves

This experiment used those PM-infected cucumber leaves that were the positive controls in the survival tests described above, i.e. pycnidia of RS1-a, C10 and G4 were present in abundance in PM colonies at the end of the experiment. To model leaf fall, the PM-infected parts of the leaves, bearing *Ampelomyces* pycnidia, were cut into approx. 2 cm^2^ pieces under a dissecting microscope; these were then placed on the soil in the respective pots that still contained the original plant. Pots continued to be kept in isolation as described above, and watered regularly, as before. Every three to six days two to four leaf pieces were removed from each pot and microscopically examined for the presence of *Ampelomyces* structures, including GFP expressing pycnidia on leaves treated with C10 and G4, as detailed above. Together with the decomposing leaf tissue samples, 0.5-1 g soil was also collected under and around the removed leaf pieces, suspended in water or 10% glycerol droplets on microscope slides, and examined under a microscope as described above. Mounting samples in water or glycerol was important as it was noted that the fluorescent signal was much weaker when samples were checked without hydration. The experiment was carried out twice.

### 2.14. Testing survival and potential development of *Ampelomyces* in autoclaved soil

Glass plates, 9 cm diam, were filled with commercial gardening soil, 10 g per plate, closed, and autoclaved twice. Conidial suspensions of the wild type RS1-a and transformants B3 and C10 were prepared and checked for viability/GFP expression on 1.5% water agar medium as detailed above. Three plates with autoclaved gardening soil were inoculated each with 300 μl of conidial suspension of one of the fungal materials by pipetting three times 100 μl of inoculum into three spots of plates that were then sealed with Parafilm, and incubated at 22°C in dark for 30 days. Three plates served as negative controls, being treated each with 300 μl sterile water. The autoclaved soil remained wet in plates sealed with Parafilm until the end of the experiment. Approx. 0.5-1 g soil samples were collected at random in each plate from ten places, seven and then 30 days after the treatments, suspended in water or 10% glycerol droplets on microscope slides, and microscopically examined for the presence of *Ampelomyces* structures, including GFP expressing structures in the case of treatments with B3 and C10, as described above. The experiment was carried out twice.

## 3. Results

### 3.1. Production and molecular characterization of transformants

Two *Ampelomyces* strains, RS1-a and GYER, isolated from rose and hornbeam powdery mildews, *Podosphaera pannosa* and *Erysiphe arcuata*, respectively, were selected for this work based on previous data on their cultural characteristics and mycoparasitic activities^31^. The two *Ampelomyces* strains are phylogenetically distantly related (Fig. 1). As a result of the *Agrobacterium tumefaciens*-mediated transformation (ATMT), hygromycin B resistant colonies appeared eight to ten weeks after co-cultivation. These all exhibited continuous, albeit slow growth in these plates, similar to the growth rate of wild type *Ampelomyces* strains^31,38^. The transformation ratio was approx. 10% for RS1-a, and 30% for GYER. Hyphae of all but one of the emerged colonies exhibited strong green fluorescence when excited with blue light (Fig. 2a); the colony that barely showed any fluorescence was excluded from further studies. Those colonies that sporulated on the selective medium produced fluorescent conidia (Fig. 2b). Colonies that were fluorescent under blue light were considered as putative GFP transformants of the respective *Ampelomyces* strain. Of these, seven were selected for RS1-a and six for GYER, and included in subsequent works. Their designations are shown in Supplementary Table S1. Their colonies were subcultured every 4-6 weeks on Czapek-Dox Agar supplemented with 2% malt (MCzA) without hygromycin B, the selection agent. Maintenance of hygromycin B resistance and GFP fluorescence in the absence of selection pressure throughout our 3-year project indicated high mitotic stability of the transgenes.

**Figure 2.**
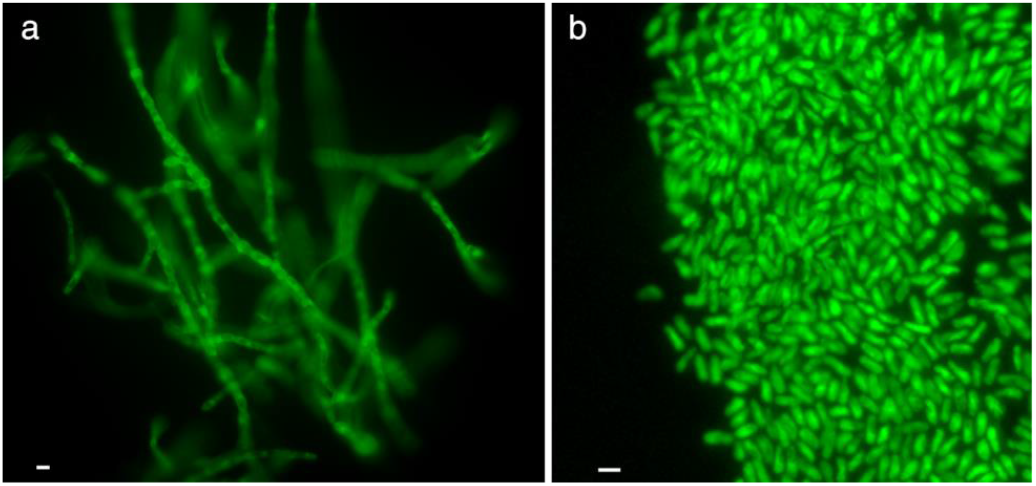
Green fluorescence emitting hyphae and conidia of transformant B3 of *Ampelomyces* strain RS1-a two weeks after its first transfer to a plate without the selection agent. (**a**) Hyphae; bar = 5 μm. (**b**) Conidia; bar = 10 μm.

PCRs targeting the amplification of *hph* and *SGFP*, two fragments of T-DNA, resulted in products with expected sizes in all the tested transformants. Amplicon sequences were identical to partial *hph* and *SGFP* fragments of pCBCT^50^, the source of T-DNA. No products were amplified from the wild type strains RS1-a and GYER during these PCRs. qPCR measurements of T-DNA copy numbers showed that eight of the selected transformants contained a single copy of the insert, four transformants had two, and one had three copies of the T-DNA (Supplementary Table S1).

### 3.2. Morphology and growth of transformants in culture

Neither morphology of coeval colonies of the seven monoconidial RS1-a transformants nor morphology of their pycnidia and conidia produced in culture differed from the characteristics of the wild type RS1-a strain. Also, colony morphology of the six GYER transformants was similar to that of the wild type strain. Pycnidia were only sporadically produced by both the wild type and the transformant GYER colonies in culture; when this happened, no differences were observed in the morphology of pycnidia and conidia of the wild type strain and the transformants. Colony growth analysis revealed significant differences in saprobic growth of four out of six GYER (p≤0.0001) and one out of seven RS1-a (p=0.004) transformants compared to the growth of the respective wild type strains (Fig. 3), but the overall colony growth patterns described for a number of genetically different *Ampelomyces* strains^31,38,51^ were not markedly altered in any of the RS1-a or GYER transformants. Two transformants with significantly higher growth rates contained one, another had two, and the fourth had three copies of T-DNA, while the other transformants characterized by growth rates similar to the wild type strain had one or two T-DNA inserts (Supplementary Table S1).

**Figure 3.**
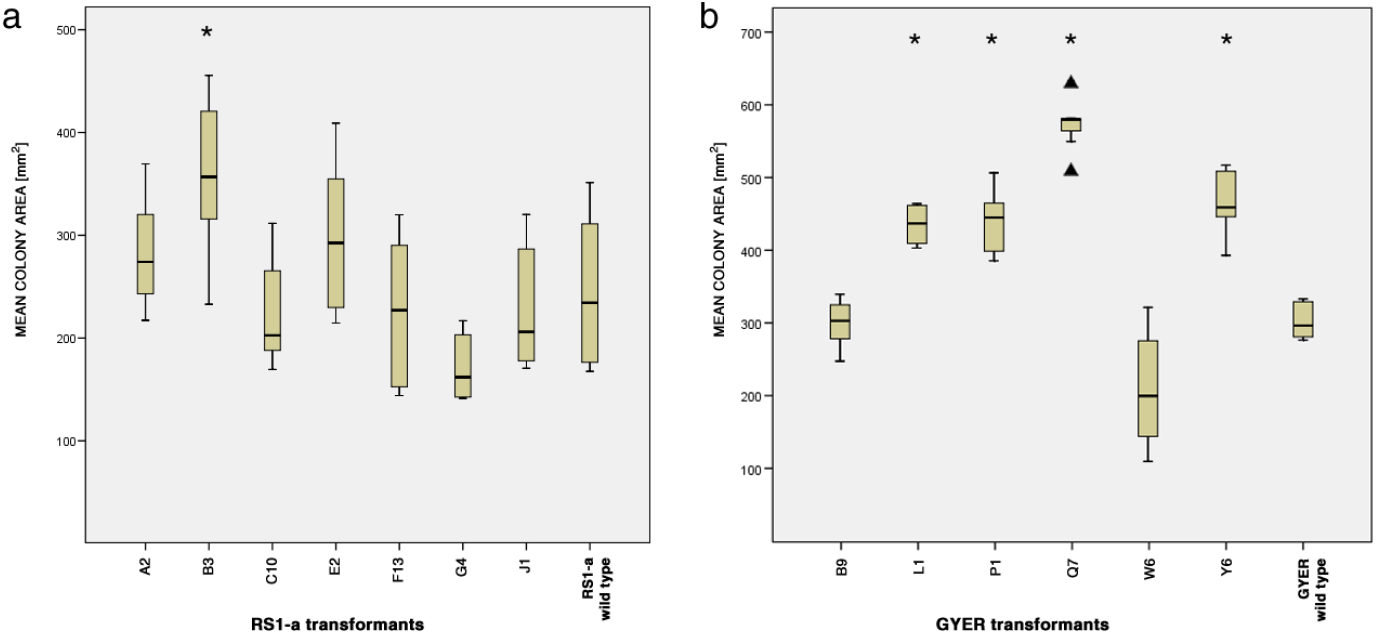
Box plot showing surface area sizes of four week old colonies of RS1-a and GYER transformants, and the respective wild types. Medians and second and third quartiles are shown as boxes; upper and lower whiskers represent the maximum and the minimum values of the dataset, respectively. Transformants exhibiting growth that was significantly different from the wild type were marked with asterisks (*; p<0.05). Triangles denote outliers. (**a**) Seven RS1-a transformants and the wild type strain. (**b**) Six GYER transformants and the wild type strain.

### 3.3. Mycoparasitic activities of transformants in mycelia of different PM species

All experiments targeting the mycoparasitic activities of transformants in PM colonies, compared to the wild type strain, were carried out with RS1-a materials. The sporulation of the wild type GYER strain and its transformants was unreliable in culture, therefore conidial suspensions could not always be prepared for mycoparasitic tests with GYER materials.

The first mycoparasitic tests were carried out with cucumber plants infected with *Podosphaera xanthii*, a common PM species infecting a number of vegetables, ornamentals and wild plants. Conidial suspensions of the seven RS1-a transformants, and the wild-type strain, were sprayed each on three PM-infected plants, and mycoparasitic interactions were examined after a 10-day incubation period. Each treatment with *Ampelomyces* resulted in development of pycnidia inside conidiophores of *P. xanthii*. Hyphae and pycnidia of all transformants, produced intracellularly in PM colonies, similar to their conidia released from intracellular pycnidia, exhibited the GFP signal when examined with a fluorescence microscope (Figs. 4a-c). Green fluorescence was not observed when hyphae, pycnidia and conidia of *Ampelomyces* were examined in samples taken from PM colonies inoculated with the wild type RS1-a strain. *Ampelomyces* pycnidia were uniformly distributed in all PM colonies on all plants. Intracellular hyphae of the transformants that entered PM conidia were much better visualized with fluorescence microscopy compared to brightfield, phase contrast and even differential interference contrast (DIC) optics (Fig. 1d). Microscopic observations did not reveal any differences amongst mycoparasitic activities of the seven transformants and the wild type strain in terms of penetration of PM hyphae, intrahyphal growth, intracellular pycnidial development inside PM conidiophores, and release of their conidia from intracellular pycnidia. No *Ampelomyces* structures were observed in samples taken from negative controls. The experiment was first carried out four, then five months after the production of transformants in culture, and then for the third time with transformants that have already been subcultured for 30 months on MCzA, without hygromycin B, the selection agent. GFP expression of hyphae, pycnidia and conidia of transformants, produced intracellularly in PM colonies, was as intensive in the third experiment as in the first two, which supported the stability of the GFP transformation in all the seven RS1-a transformants. This was also shown by GFP signal emission and resistance to 50 mg/l hygromycin B of transformant B3 in culture, after being re-isolated from a parasitized *P. xanthii* colony.

**Figure 4.**
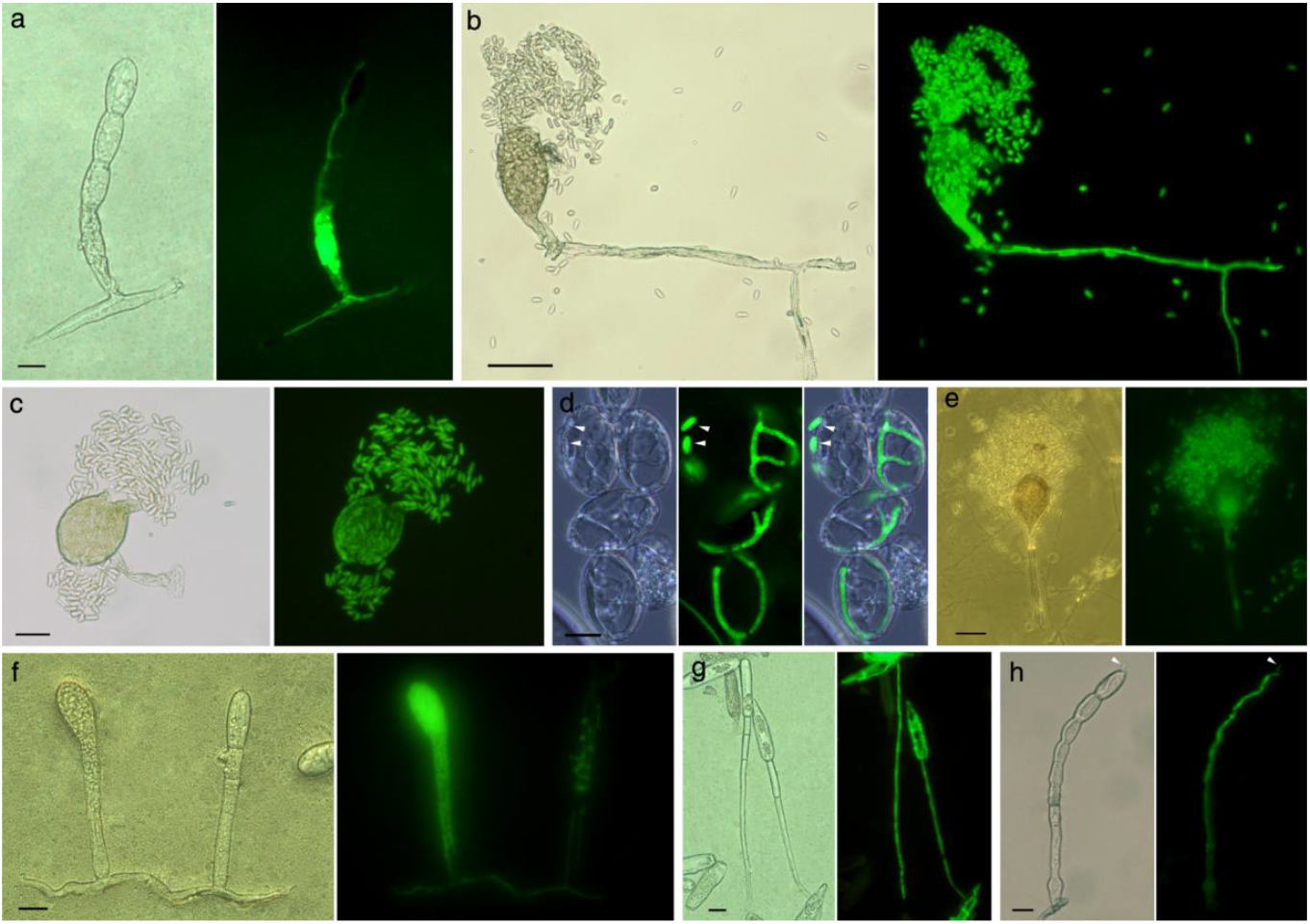
Mycoparasitism of GFP-expressing transformants of *Ampelomyces* strain RS1-a in five powdery mildew species, on their host plants, 10 days post inoculations. (**a**) A young pycnidium of transformant J1 developing in the second cell of a conidiophore of *Podosphaera xanthii* on cucumber. Note also intracellular hyphae colonizing all the other cells of the conidiophore. Left: micrograph with DIC optics; right: fluorescence microscopy. Bar = 20 μm. (**b**) A mature pycnidium of transformant G4 developed in a conidiophore of *P. xanthii* releasing conidia following rupture of the intracellular pycnidial wall. Note also intracellular hyphae in other parts of the mycelium. Left: DIC optics; right: fluorescence microscopy. Bar = 50 μm. (**c**) A mature pycnidium of transformant B3 developed in *P. xanthii* and releasing conidia. Left: DIC optics; right: fluorescence microscopy. Bar = 50 μm. (**d**) A close-up of hyphae of transformant G4 in conidia of *P. xanthii*. White arrows point to *Ampelomyces* conidia, outside of *P. xanthii* conidia. Left to right: DIC optics, fluorescence microscopy, and overlap of these two images. Bar = 10 μm. (**e**) A pycnidium of transformant B3 developed in a conidiophore of *Erysiphe necator* on grape and releasing conidia. Left: bright field optics; right: fluorescence microscopy. Bar = xxx. (**f**) Two conidiophores of *Pseudoidium neolycopersici* on tomato parasitized by transformant G4. A young pycnidium is developing in the conidiophore on the left; this process is even less advanced in the conidiophore on the right, where intracellular hyphae started to anastomose in the second and third cell. Note also intracellular hyphae in other parts of the mycelium. Left: DIC optics; right: fluorescence microscopy. Bar = 20 μm. (**g**) Hyphae of transformant G4 growing inside conidiophores and conidia of *Leveillula taurica* on pepper. Left: DIC optics; right: fluorescence microscopy. Bar = 20 μm. (**h**) Hyphae of transformant C10 growing inside a conidiophore of *Blumeria graminis* f. sp. *hordei* on barley. Note that the hypha grew out of the most distal PM conidium produced on this conidiophore (white arrow). Left: DIC optics; right: fluorescence microscopy. Bar = 20 μm. Micrographs in panels a, c and e-h were taken from samples made with cellotape that reduced DIC effects but did not influence the quality of images taken with fluorescence microscopy.

Mycoparasitic tests were done in the same way with the following PMs: *Pseudoidium neolycopersici* infecting tomato, *Leveillula taurica* infecting pepper, *Blumeria graminis* infecting barley, and *Erysiphe necator* infecting grapevine. Again, visualization of intracellular structures of *Ampelomyces*, and especially the visualization of their hyphae inside the hyphae, conidiophores and conidia of the parasitized PMs, was much better with fluorescence microscopy compared to brightfield, phase contrast and DIC optics (Figs. 4e-h). Furthermore, confocal laser scanning microscopy (CLSM) revealed that the cells of the walls of transformants’ mature intracellular pycnidia did not exhibit a strong GFP signal (Fig. 5). When intracellular pycnidia were mature, with dark brown wall cells, it was mainly the green fluorescence of *Ampelomyces* conidia localized inside pycnidia that exhibited the GFP signal inside PM conidiophores under blue light. In some cases, this was also somewhat visible with fluorescence microscopy (Fig. 4b,c,e), but it was only CLSM that clearly revealed the lack of strong GFP signal from the walls of transformants’ pycnidia (Fig. 5).

**Figure 5.**
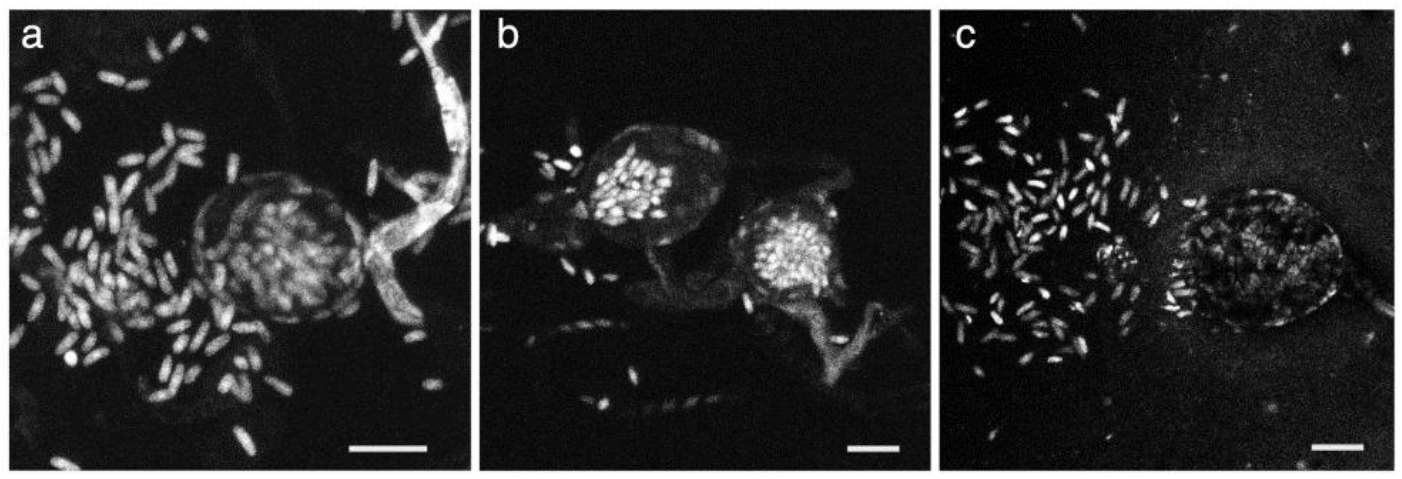
Pycnidia of transformant J1 of *Ampelomyces* strain RS1-a developed inside conidiophores of *Blumeria graminis* f. sp. *hordei* on barley, 10 days post inoculations. Samples were observed with confocal laser scanning microscope (CLSM) and images of optical slices were used to prepare z-stacked projections shown here. (**a**) A mature pycnidium produced in the foot-cell of a conidiophore and releasing conidia. Bar = 20 μm. (**b**) Two intracellular pycnidia; release of conidia was observed from the one on the left. Bar = 20 μm. (**c**) An intracellular pycnidium that has already released most of its conidia. Bar = 20 μm. All images show that the GFP signal is very weak, or cannot be detected, from the cells of the intracellular pycnidial walls, while GFP emission of conidia is always strong.

### 3.4. Quantification of mycoparasitic activity of transformants in cucumber PM colonies

Mycoparasitic activity of the seven transformants and the wild type strain was quantified based on the number of pycnidia produced in the conidiophores of *P. xanthii* on cucumber leaves, determined using an already established method^31,40^. No significant differences (p>0.1) were found amongst the values determined for the individual treatments with the seven transformants compared to the wild type (Supplementary Table S1), the average number of intracellular pycnidia being 811 ± 66 pycnidia per cm^2^ parasitized PM mycelium for transformants and 848 ± 178 pycnidia per cm^2^ PM mycelium for the wild type strain (Fig. 6).

**Figure 6.**
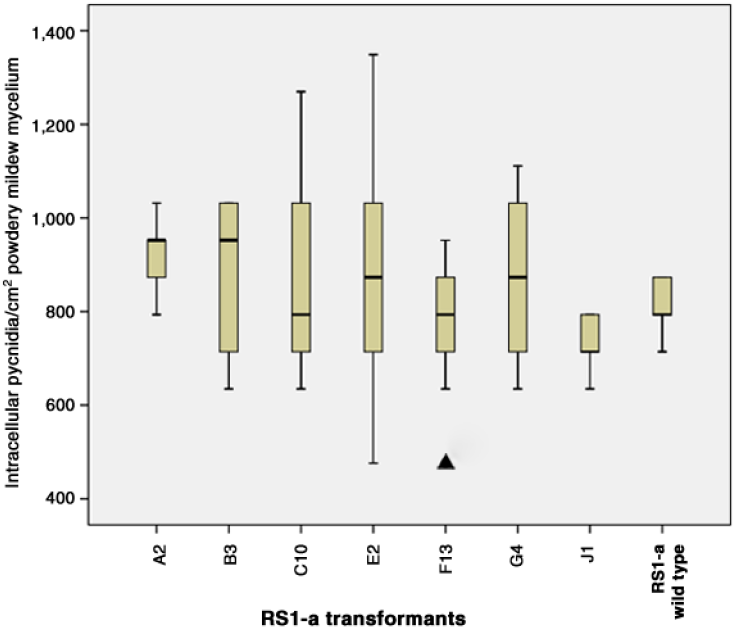
Box plot of mycoparasitic activities of seven RS1-a transformants and the wild type expressed as number of intracellular pycnidia developed in conidiophores of *Podosphaera xanthii* on cucumber per cm^2^ powdery mildew mycelium. Medians and second and third quartiles are shown as boxes; upper and lower whiskers represent the maximum and the minimum values of the dataset, respectively. No significant differences from RS1-a wild type were detected (p>0.1). Triangle denotes outlier.

### 3.5. Testing survival and potential development of *Ampelomyces* on cucumber plants in the absence of the mycohost

To find out whether *Ampelomyces* conidia can survive for at least 2-3 weeks on PM-free host plant surfaces, and if so, whether such survived conidia can still initiate mycoparasitism of newly established PM colonies on the same host plant surfaces, two transformants, C10 and G4, and the wild type RS1-a strain were sprayed each on PM-free cucumber plants. Ten days following treatments fluorescent microscopy revealed that conidia of the transformants germinated and the emerged hyphae started to grow on the leaves (Supplementary Fig. S1). This was also detected with brightfield microscopy on plants inoculated with the wild type strain. The visualization of the transformants’ conidia with fluorescence microscopy was clearly superior to the detection of the germinated wild type *Ampelomyces* conidia on leaf samples. No saprobic pycnidial production was observed on any leaves in the absence of PM.

Four, seven, 10, 12, 14, 18 and 21 days after being sprayed with one of the two transformants, or the wild type strain, or water, six cucumber plants per treatment were infected with *P. xanthii*. Each infected plant cohort was examined 10 days after PM infection for the presence of *Ampelomyces* structures in the newly developed PM colonies. Mature intracellular pycnidia of *Ampelomyces*, releasing conidia, were detected in various numbers, and in small areas, of the young PM colonies developed in 10 days on the infected plants, including those that were infected with *P. xanthii* 21 days after being treated with one of the two transformants or the wild type strain. Therefore, *Ampelomyces* survived for at least three weeks on PM-free cucumber plants, and was able penetrate and parasitize newly developed PM colonies even after this survival period. However, the level of mycoparasitism of PM colonies that have started to develop on leaves ten to 21 days after *Ampelomyces* treatments was low, and the presence of *Ampelomyces* structures in PM hyphae and conidiophores was patchy, in contrast to what was seen in positive controls, and also in previous experiments, where PM colonies have already covered large leaf areas when they were sprayed with *Ampelomyces*. GFP expression of transformants has greatly enhanced the study of the respective samples compared to those taken from PM colonies developed on leaves pre-treated with the wild type strain (Fig. 7)

**Figure 7.**
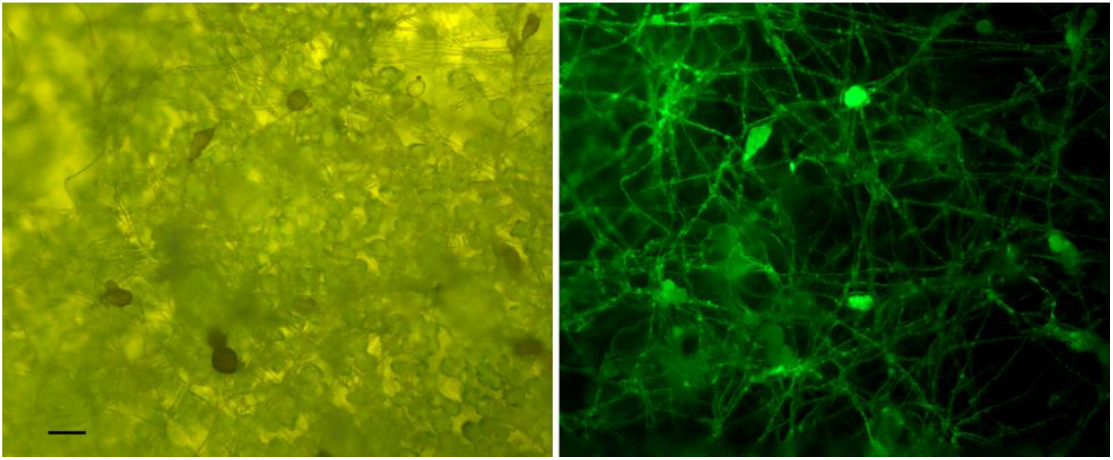
Intracellular hyphae and pycnidia of transformant B3 of *Ampelomyces* strain RS1-a developed in young, 10-day old colonies of *Podosphaera xanthii* on cucumber. Conidia of the transformant were sprayed on cucumber plants four days before the powdery mildew inoculum was dusted onto these. Left: bright field; right: fluorescence microscopy. Bar = 50 μm.

### 3.6. Testing survival and potential development of *Ampelomyces* in senescent/decomposed PM-infected leaves on the soil

*Ampelomyces* structures on PM-infected leaf pieces placed on the soil in the respective plant’s pots were detected for 30 days from the start of the experiment. Bright fluorescence of GFP transformants was observed during the first 9-12 days in the sampled, and gradually decomposing leaf pieces; in samples examined after this period the fluorescent signal was always weaker. *Ampelomyces*pycnidia found on leaf pieces have usually released conidia into the mounting medium during microscopic examinations, and not before, although the soil in pots was continuously kept wet. GFP fluorescence was not detected on leaf pieces inoculated with the wild type strain, but background fluorescence of plant material and soil particles was always observed. Newly developed *Ampelomyces* hyphae were rarely seen in the senescent leaf tissues, while saprobic production of pycnidia in the decomposing leaves, and/or the adjacent soil samples collected from the pots, was not observed in this experiment.

### 3.7. Testing survival and potential development of *Ampelomyces* in autoclaved soil

*Ampelomyces* conidia germinated and the emerging hyphae continued to grow in the sterile environment set up in this experiment. Hyphae of transformants were found on the surface of soil particles and also between particles as aerial hyphae. Wild type hyphae were only detected as aerial hyphae probably due to difficulties in observing soil samples with brightfield, phase contrast, and DIC microscopy. GFP expression of transformants greatly enhanced the visualization of hyphae on and around soil particles (Fig. 8). Pycnidia of *Ampelomyces* were not detected in any plates with autoclaved soil in this 30-day long experiment.

**Figure 8.**
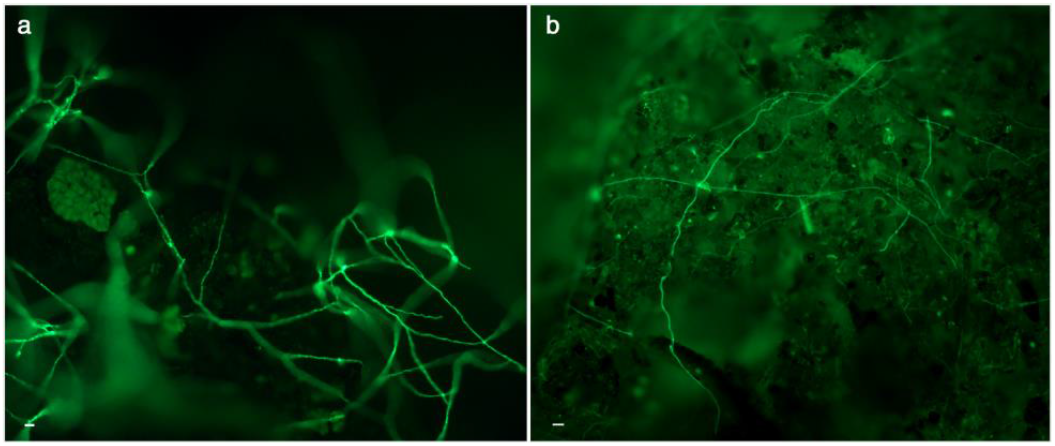
Hyphae of transformant B3 of *Ampelomyces* strain RS1-a in plates with autoclaved soil, 30 days post inoculation. (a) Aerial hyphae of the transformant. Bar = 20 μm. (b) Hyphae attached to a soil particle. Bar = 20 μm.

## 4. Discussion

Heterologous fluorescent protein expression has been used in the study of mycoparasitic interactions, enhancing the visualization of interfungal parasitic relationships in diverse settings^15–19^. This technology has been proven extremely valuable in the study of the interactions between PMs and *Ampelomyces*, for several reasons. Firstly, the detection of *Ampelomyces* structures, and especially hyphae, was easier *per se* when diverse samples containing GFP transformants were examined with fluorescence microscopy compared to brightfield, phase contrast and/or DIC observations of the same samples, or samples containing wild type *Ampelomyces* structures. Intracellular *Ampelomyces* hyphae have been identified with brightfield microscopy since the 19^th^ century, first by De Bary^23^, and then also with other optics during their development inside PM hyphae, conidiophores and conidia^25,28,48^. However, detection of *Ampelomyces* could be difficult especially during the early stages of the mycoparasitic interaction, before the production of intracellular pycnidia in PM conidiophores. GFP expression of transformants has greatly enhanced the visualization of their hyphae right after conidial germination on PM-free host plant surfaces, soon after their penetration into powdery mildew hyphae, and also on the surface of soil particles, and distinguished these from hyphae of other fungi in experiments carried out in non-sterile environments.

Secondly, the improved visualization of *Ampelomyces* structures made it possible to perform experiments that focused on deciphering the controversial environmental fate of *Ampelomyces*^44,45^ before and after its well-known mycoparasitic relationship with PM colonies. Inoculations of PM-free cucumber leaves with *Ampelomyces* revealed that these mycoparasites can persist up to 21 days on host plant surfaces without being in contact with their mycohosts. After this time frame, *Ampelomyces* was still able to initiate mycoparasitism, i.e. penetrate PM structures that were provided experimentally on these plants. This has not been previously shown, and the general assumption was that actively growing PM colonies are always needed for the establishment of *Ampelomyces*^22,24,30,32^. This result is also important from a biocontrol perspective because it indicates that commercial *Ampelomyces* biofungicide products sprayed on those parts of the treated crops that are not infected with PM at the time of the application may not be completely wasted. Those amounts of biofungicide may still have an impact on controlling PM infections, in a preventive way, because they can initiate mycoparasitism of PM colonies that may appear later on the respective plant surfaces. However, our results have also indicated that a part of the *Ampelomyces* inoculum was lost during the up to 21-day long PM-free period on host plant surfaces. After the establishment of PM colonies on these plants, the level of mycoparasitism was much lower and patchy compared to plants that were treated with *Ampelomyces* when large leaf areas were already covered with PM, and the mycoparasites could readily attack PM colonies. This was not surprising as the 10-day old PM colonies developed on those experimental plants were too young to support more mycoparasitism.

Two experiments were carried out to decipher what happens to *Ampelomyces* on fallen senescent and decomposing PM-infected leaves on the ground, and in the soil, during the post-mycoparasitic phase of their life cycle. When PM-infected leaf pieces bearing transformants’ pycnidia were placed on the soil in pots, in a non-sterile environment, fluorescent microscopy revealed that the GFP signal in samples started to decrease after 10 days, and disappeared after 30 days. This was interpreted as a reduction, and the end of the GFP gene expression, i.e. transformants’ active metabolism, respectively. Most importantly, saprobic production of new pycnidia, thus new sporulation, was never observed either in the decomposing leaf tissues and adjacent soil samples, or in the second experiment that consisted of inoculations of autoclaved soil volumes with *Ampelomyces*. The autoclaved soil remained wet until the end of the 30-day experiment; this was regarded as a favourable condition for saprobic pycnidial production. Pycnidia of *Ampelomyces* were not detected on PM-free leaves either, during our experiment targeting the persistence of these mycoparasites on host plant surfaces in the absence of their mycohosts. In fact, in this study, saprobic development of *Ampelomyces* pycnidia was observed only in culture, similar to earlier works^9,29,32,39,41,42,51,52^. According to this study, saprobic development of *Ampelomyces* prior to the mycoparasitic phase of its life cycle was limited to hyphal growth on PM-free plant surfaces; and to local metabolic activity, without any signs of extensive new hyphal growth and new pycnidial production, in/around decomposing PM-infected leaf tissues during the post-mycoparasitic phase. These results indicated that *Ampelomyces*, similar to PMs, occupies a niche in the phyllosphere, and acts there primarily as an intracellular mycoparasite of PMs. Moreover, our results showed that *Ampelomyces*was not able to sporulate in the absence of its mycohosts, in either the phyllosphere or the soil level. It may persist in the environment without its mycohosts, but based on these results it is unlikely that it is able to colonize niches other than PM colonies. Therefore, as suggested earlier^45^, concerns regarding the presumed impact of *Ampelomyces* on soil microbiota^44^ remain unconfirmed. Overwintering of *Ampelomyces* as saprobically produced pycnidia in the leaf debris, hypothesized in the early mycological literature^23,49^, has never been confirmed either. All experimental studies on overwintering forms of *Ampelomyces* have revealed that these were always linked to PM structures^29,40,48,53^.

To develop *Ampelomyces* strains constitutively expressing GFP we utilized ATMT, ‘a silver bullet in a golden age of functional genomics’^54^. The transformation ratios were similar to those achieved in other studies^55^. GFP fluorescence and hygromycin B resistance of transformants indicated that the heterologous genes and regulator regions provided on the pCBCT plasmid were functional in both *Ampelomyces* strains. These transformants have maintained their hygromycin B resistance and GFP fluorescence for almost three years of subculturing in absence of selection pressure, being mitotically stable. Pycnidia of transformants developed in culture and also in PM conidiophores produced numerous conidia; these were also fluorescent. This has also indicated the mitotic stability of the integrated transgenes. Cells of the pycnidial walls of *Ampelomyces* were the only structures in transformants that did not exhibit strong GFP signal, as revealed by CLSM.

All the mycoparasitic tests showed that transformants penetrated PM colonies, and sporulated inside PM conidiophores similar to the wild type strain. Quantitative data revealed that the level of mycoparasitism did not differ in transformants and the wild type strain. This indicated that none of the genes that play a role in mycoparasitism were disrupted during GFP transformation. Significant differences (p≤0.004) were, however, detected in the growth rate of some transformants in culture, compared to the wild type strain. Therefore, transformation may have still had an identifiable impact on the target organisms. Most transformants had a single copy of the T-DNA in their genome; the significant differences in colony growth rate could not be linked to the number of T-DNA copies in different transformants. Further studies can utilize those transformants that had a single T-DNA copy and did not exhibit any significant differences in their cultural characteristics when compared to the wild type strain.

The transformation method was equally successful in two phylogenetically distinct *Ampelomyces* strains that were markedly different from a physiological point of view, as well: RS1-a has always intensively sporulated in culture, and has already been selected as a potential BCA^31^, while GYER was characterized by poor and only occasional sporulation in culture. This may indicate that ATMT could be used to transform phylogenetically and physiologically diverse *Ampelomyces* strains. Such transformations may target functional genomics analyses of genes involved in mycoparasitism. A transcriptome analysis in *Ampelomyces*^35^ has already provided a useful database for such studies. Therefore, the present work, in addition to shedding light on largely unknown aspects of the life cycle of a widespread mycoparasite, has also established the framework for a molecular genetic toolbox to be used in future studies on *Ampelomyces*.

### Data availability

All data generated or analysed during this study are included in this manuscript (and its Supplementary Tables). Newly determined DNA sequences were deposited in NCBI GenBank under accession numbers MH879020-MH879022. There are no restrictions on the availability of any data supporting this study.

## Acknowledgements

This work was supported by the Széchenyi 2020 programme, the European Regional Development Fund and the Hungarian Government (GINOP-2.3.2-15-2016-00061). A grant of the Hungarian Research, Development and Innovation Office (NKFIH NN100415) and a grant of the Austrian-Hungarian Action Foundation (90öu16) have also supported a part of this work. Alexandra Pintye’s contribution was supported by János Bolyai Research Scholarship of the Hungarian Academy of Sciences (MTA). The authors are grateful to Csilla Rohm, Ildikó Csorba and Dr. Endre Tóth for their technical assistance and to Dr. Klára Mészáros for providing the barley genotype used in this work.

## Author contributions

LK, MG and GMK conceived, designed and coordinated the research. MZN performed the GFP transformation work and the characterization of the transformants. MZN and AP performed the mycoparasitic tests and DIC and fluorescence microscopy work, PV the CLSM study, ÁNH the statistical tests. MZN, LK and MG wrote the paper and all the authors commented on the paper.

## Additional Information

Supplementary information accompanies this paper. This consists of one figure and four tables.

## SUPPLEMENTARY FIGURE

**Supplementary Figure S1.**
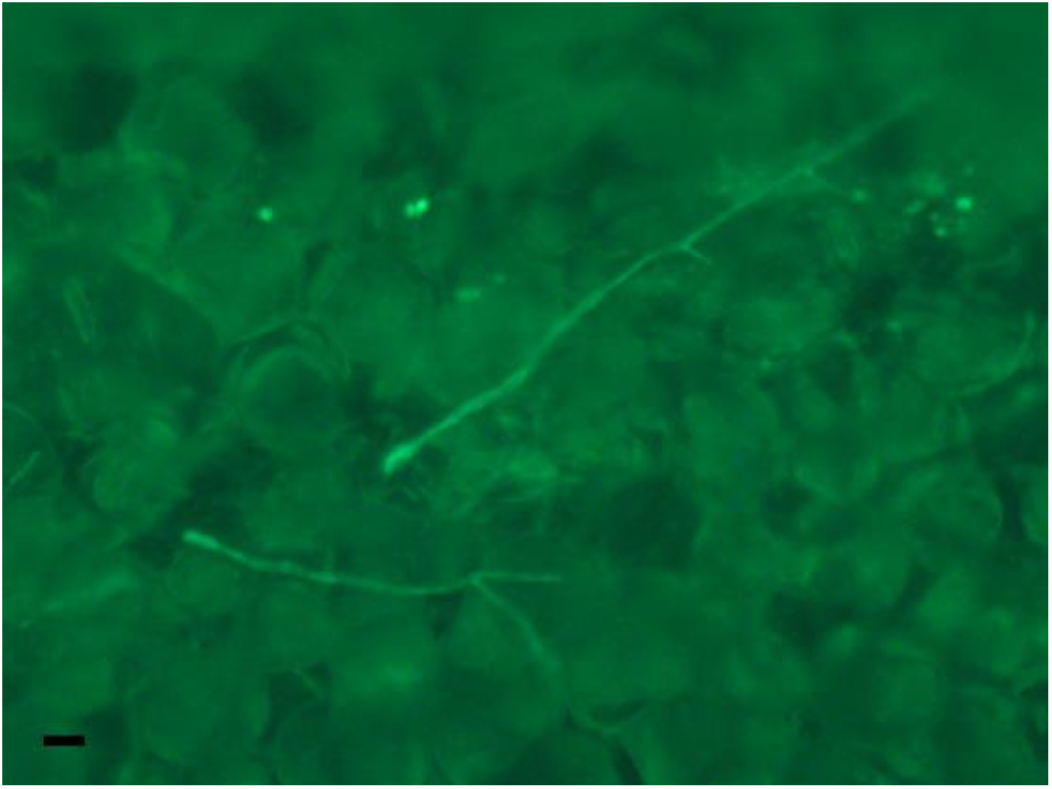
Conidia of transformant B3 of *Ampelomyces* strain RS1-a germinated on powdery mildew-free cucumber leaves. The micrograph was taken ten days after inoculation. Bar = 10 μm.

## SUPPLEMENTARY TABLES

**Supplementary Table S1.** Designations of the six selected transformants of *Ampelomyces* strain GYER, and seven selected transformants of strain RS1-a, and their characteristics in terms of T-DNA copy numbers, growth in culture, and mycoparasitic activity in *Podosphaera xanthii* on cucumber.

| Transformant designation | T-DNA copy number | Growth in culture compared to wild type, and corresponding p values |  | Mycoparasitic activity in *P. xanthii* compared to wild type, and corresponding p values |  |
| --- | --- | --- | --- | --- | --- |
| Strain GYER |  |  |  |  |  |
| B9 | 2 | No difference | 1.000 | ND* | - |
| L1 | 1 | Faster | 0.000001 | ND | - |
| P1 | ND | Faster | 0.0001 | ND | - |
| Q7 | 1 | Faster | 0.000000004 | ND | - |
| W6 | 1 | No difference | 0.132 | ND | - |
| Y6 | 2 | Faster | 0.0002 | ND | - |
| Strain RS1-a |  |  |  |  |  |
| A2 | 1 | No difference | 0.746 | No difference | 0.228 |
| B3 | 3 | Faster | 0.004 | No difference | 0.988 |
| C10 | 1 | No difference | 0.978 | No difference | 0.999 |
| E2 | 1 | No difference | 0.423 | No difference | 0.999 |
| F13 | 1 | No difference | 0.984 | No difference | 0.999 |
| G4 | 2 | No difference | 0.120 | No difference | 1.000 |
| J1 | 2 | No difference | 0.994 | No difference | 0.196 |

\*ND = not determined

**Supplementary Table S2.**
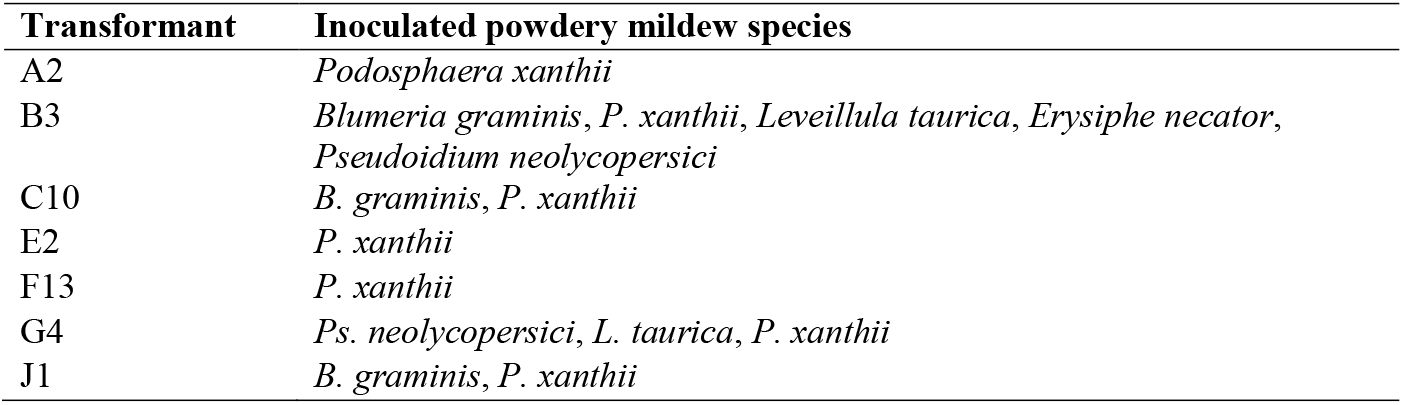
List of transformants of *Ampelomyces* strain RS1-a used in the mycoparasitic tests carried out with five powdery mildew species on their respective host plants.

**Supplementary Table S3.**
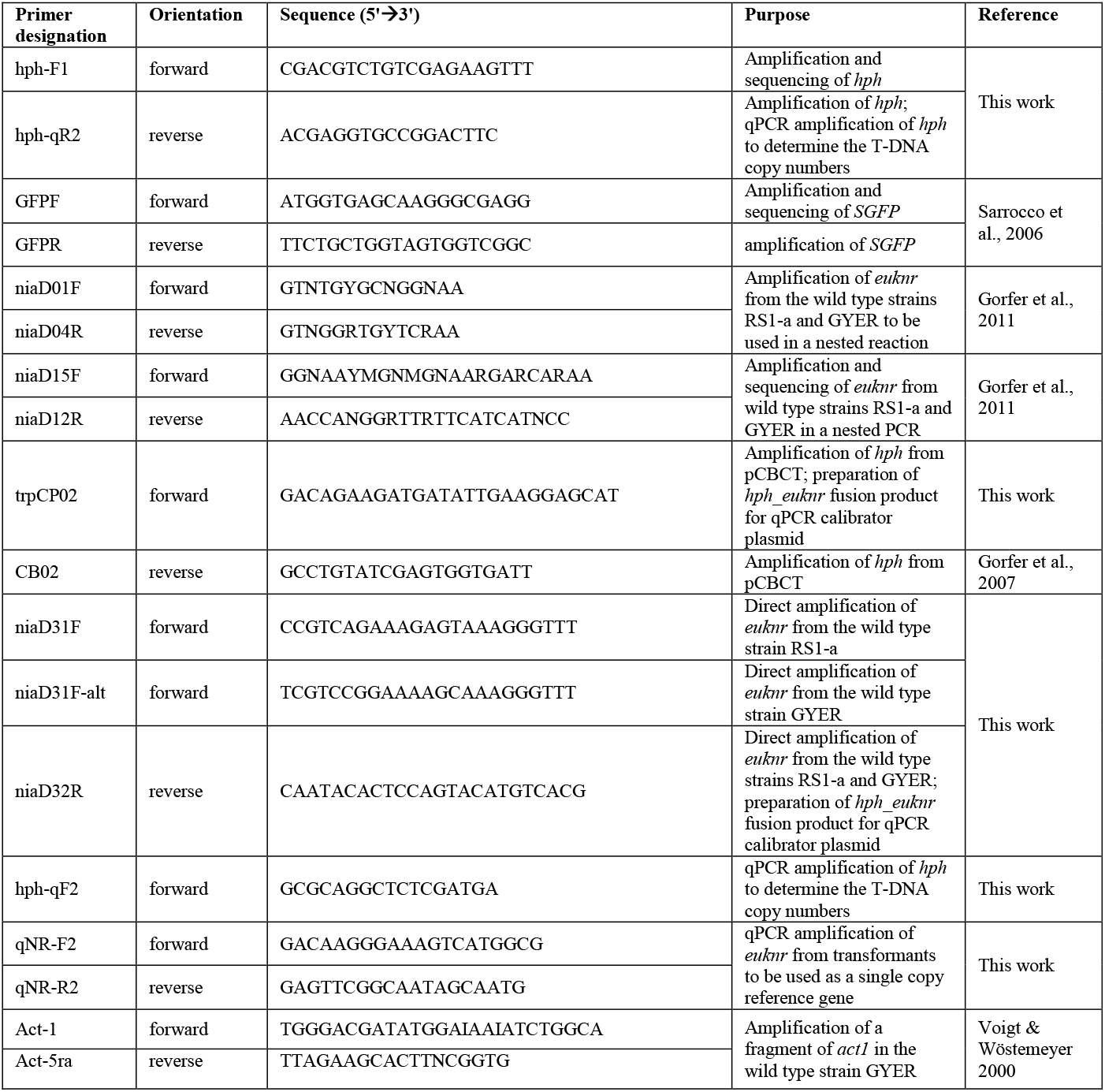
Primers used in this work.

**Supplementary Table S4.** Descriptive statistics, Levene’s Test of Equality of Error Variances and Tests of Normality of the three datasets examined.

| Descriptive Statistics - Mycoparasitism |  |  |  | Descriptive Statistics - GYER transformants' growth |  |  |  | Descriptive Statistics - RS1-a transformants' growth |  |  |  |
| --- | --- | --- | --- | --- | --- | --- | --- | --- | --- | --- | --- |
| Transformant | Mean | Std. Deviation | N | Transformant | Mean | Std. Deviation | N | Transformant | Mean | Std. Deviation | N |
| A2 | 917.108 | 89.714 | 9 | Y6 | 468.320 | 45.941 | 7 | A2 | 282.665 | 51.767 | 8 |
| B3 | 881.836 | 155.950 | 9 | B9 | 299.831 | 31.521 | 8 | B3 | 359.415 | 75.190 | 8 |
| C10 | 890.653 | 227.190 | 9 | L1 | 435.372 | 26.719 | 8 | C10 | 224.156 | 50.232 | 8 |
| E2 | 899.473 | 266.199 | 9 | P1 | 438.591 | 42.280 | 8 | E2 | 297.260 | 76.026 | 8 |
| F13 | 758.378 | 143.688 | 9 | Q7 | 573.843 | 34.467 | 8 | F13 | 225.435 | 74.868 | 8 |
| G4 | 864.200 | 165.740 | 9 | W6 | 208.580 | 76.876 | 8 | G4 | 171.675 | 32.152 | 8 |
| J1 | 723.105 | 62.042 | 9 | GYER wild type | 302.796 | 24.180 | 8 | J1 | 228.953 | 61.183 | 8 |
| RS1-a wild type | 811.290 | 66.138 | 9 | Total | 388.188 | 123.872 | 55 | RS1-a wild type | 245.322 | 73.366 | 8 |
| Total | 843.255 | 168.194 | 72 |  |  |  |  | Total | 254.360 | 80.744 | 64 |
| Levene's Test of Equality of Error Variances |  |  |  | Levene's Test of Equality of Error Variances |  |  |  | Levene's Test of Equality of Error Variances |  |  |  |
| F | df1 | df2 | Sig. | F | df1 | df2 | Sig. | F | df1 | df2 | Sig. |
| 4.679 | 7 | 64 | 0.000 | 3.875 | 6 | 48 | 0.003 | 1.488 | 7 | 56 | 0.190 |
| Tests of Normality |  |  |  | Tests of Normality |  |  |  | Tests of Normality |  |  | Skewness |
| Kolmogorov-Smirnov |  |  |  | Kolmogorov-Smirnov |  |  |  | Kolmogorov-Smirnov |  |  | 0.299 |
| Statistic | df | Sig. |  | Statistic | df | Sig. |  | Statistic | df | Sig. | Kurtosis |
| 0.074 | 72 | 0.200* |  | 0.067 | 55 | 0.200* |  | 0.110 | 64 | 0.051 | 0.590 |

## References

1 Jeffries, P. & Young, T. W. Interfungal Parasitic Relationships. (CAB International, Wallingford, UK, 1994).

2 Kiss, L. The role of hyperparasites in host plant-parasitic fungi relationships. In Biotic Interactions in PlantPathogen Associations (eds M. J. Jeger & N. J. Spence), 227–236 (CAB International, Wallingford, UK, 2001).

3 Viterbo, A. & Horwitz, B. A. Mycoparasitism. In Cellular and Molecular Biology of Filamentous Fungi (eds Borkovich K. A. & Ebbole D. J.), 676–693 (ASM Press, Washington, D.C., 2010).

4 Boddy, L. Interactions between fungi and other microbes. In The Fungi (Third Edition) (eds Watkinson S. C., Boddy L., & Money N. P.), 337–360 (Elsevier, 2015).

5 Tollenaere, C. et al. A hyperparasite affects the population dynamics of a wild plant pathogen. Mol. Ecol. 23, 5877–5887 (2014). doi:10.1111/mec.12908

6 Parratt, S. R., Barres, B., Penczykowski, R. M. & Laine, A. L. Local adaptation at higher trophic levels: contrasting hyperparasite-pathogen infection dynamics in the field and laboratory. Mol. Ecol. 26, 1964–1979 (2017). doi:10.1111/mec.13928

7 Parratt, S. R. & Laine, A.-L. Pathogen dynamics under both bottom-up host resistance and top-down hyperparasite attack. J. Appl. Ecol. (2018). doi:10.1111/1365-2664.13185

8 Parratt, S. R. & Laine, A. L. The role of hyperparasitism in microbial pathogen ecology and evolution. ISME J. 10, 1815–1822 (2016). doi:10.1038/ismej.2015.247

9 Liyanage, K. K. et al. Morpho-molecular characterization of two *Ampelomyces* spp. (Pleosporales) strains mycoparasites of powdery mildew of *Hevea brasiliensis*. Front. Microbiol. 9, 1–10 (2018). doi:10.3389/fmicb.2018.00012

10 Calonje, M., Mendoza, C. G., Cabo, A. P., Bernardo, D. & Novaes-Ledieu, M. Interaction between the mycoparasite *Verticillium fungicola* and the vegetative mycelial phase of *Agaricus bisporus*. Mycol. Res. 104, 988–992 (2000).

11 Jacobs, K., Holtzman, K. & Seifert, K. A. Morphology, phylogeny and biology of *Gliocephalis hyalina*, a biotrophic contact mycoparasite of *Fusarium* species. Mycologia 97, 111–120 (2005).

12 Smith, S., Prince, M. & Whipps, J. Characterization of *Sclerotinia* and mycoparasite *Coniothyrium minitans* interaction by microscale co-culture. Lett. Appl. Microbiol. 47, 128–133 (2008).

13 Kim, S. H. & Vujanovic, V. Changes in mycoparasite-*Fusarium* hosts interfaces in response to hostile environment as revealed by water contact angle and atomic force microscopy. Biol. Control 121, 247–255 (2018).

14 Lorang, J. M. et al. Green fluorescent protein is lighting up fungal biology. Appl. Environ. Microbiol. 67, 1987–1994 (2001). doi:10.1 128/AEM.67.5.1987-1994.2001

15 Lübeck, M. et al. GUS and GFP transformation of the biocontrol strain *Clonostachys rosea* IK726 and the use of these marker genes in ecological studies. Mycol. Res. 106, 815–826 (2002). doi:10.1017/s095375620200607x

16 Lu, Z. et al. In vivo study of *Trichoderma*-pathogen-plant interactions, using constitutive and inducible green fluorescent protein reporter systems. Appl. Environ. Microbiol. 70, 3073–3081(2004). doi:10.1128/aem.70.5.3073-3081.2004

17 Sarrocco, S. et al. Histopathological studies of sclerotia of phytopathogenic fungi parasitized by a GFP transformed *Trichoderma virens* antagonistic strain. Mycol. Res. 110, 179–187 (2006).

18 Bitsadze, N., Siebold, M., Koopmann, B. & Tiedemann, A. Single and combined colonization of *Sclerotinia sclerotiorum* sclerotia by the fungal mycoparasites *Coniothyrium minitans* and *Microsphaeropsis ochracea*. Plant Pathol. 64, 690–700 (2015).

19 Amey, R. C. et al. PEG-mediated and *Agrobacterium*-mediated transformation in the mycopathogen *Verticillium fungicola*. Mycol. Res. 106, 4–11 (2002).

20 Glawe, D. A. The powdery mildews: a review of the world’s most familiar (yet poorly known) plant pathogens. Annu. Rev. Phytopathol. 46, 27–51 (2008). doi:10.1146/annurev.phyto.46.081407.104740

21 Braun, U. & Cook, R. T. A. Taxonomic Manual of the Erysiphales (Powdery Mildews). (CBS-KNAW Fungal Biodiversity Centre, Utrecht, The Netherlands, 2012).

22 Kiss, L. Intracellular mycoparasites in action: Interactions between powdery mildew fungi and *Ampelomyces*. In Stress in Yeasts and Filamentous Fungi (eds Avery S. V., Stratford M., & Van West P.), 37–52 (Academic Press, London, UK, 2008).

23 De Bary, A. *Eurotium, Erysiphe, Cicinnobolus*, nebst Bemerkungen über die Geschlechtsorgane der Ascomyceten. In Beiträge zur Morphologie und Physiologie der Pilze (eds De Bary A. & Woronin M.), 1–95 (Winter Verlag, Frankfurt A.M., Germany, 1870).

24 Kiss, L., Russell, J. C., Szentivãnyi, O., Xu, X. & Jeffries, P. Biology and biocontrol potential of *Ampelomyces* mycoparasites, natural antagonists of powdery mildew fungi. Biocontrol Sci. Technol. 14, 635–651 (2004). doi:10.1080/09583150410001683600

25 Speer, E. Beitrag zur Morphologie von *Ampelomyces quisqualis* Ces. Sydowia 31, 242–246 (1979).

26 Hashioka, Y. & Nakai, Y. Ultrastructure of pycnidial development and mycoparasitism of *Ampelomyces quisqualis* parasitic on *Erysiphales*. Trans. Mycol. Soc. Japan 21, 329–338 (1980).

27 Sundheim, L. & Krekling, T. Host-parasite relationships of the hyperparasite *Ampelomyces quisqualis* and its powdery mildew host *Sphaerotheca fuliginea*. J. Phytopathol. 104, 202–210 (1982).

28 Kiss, L. et al. Microcyclic conidiogenesis in powdery mildews and its association with intracellular parasitism by *Ampelomyces*. Eur. J. Plant Pathol. 126, 445–451 (2010).

29 Falk, S. P., Gadoury, D. M., Cortesi, P., Pearson, R. C. & Seem, R. C. Parasitism of *Uncinula necator* cleistothecia by the mycoparasite *Ampelomyces quisqualis*. Phytopathology 85, 794–800 (1995).

30 Falk, S. P., Gadoury, D. M., Pearson, R. C. & Seem, R. C. Partial control of grape powdery mildew by the mycoparasite *Ampelomyces quisqualis*. Plant Dis. 79, 483–490 (1995).

31 Legler, S. E. et al. Sporulation rate in culture and mycoparasitic activity, but not mycohost specificity, are the key factors for selecting *Ampelomyces* strains for biocontrol of grapevine powdery mildew (*Erysiphe necator*). Eur. J. Plant Pathol. 144, 723–736 (2016). doi:10.1007/s10658-015-0834-1

32 Sztejnberg, A., Galper, S., Mazar, S. & Lisker, N. *Ampelomyces quisqualis* for biological and integrated control of powdery mildews in Israel. J. Phytopathol. 124, 285–295 (1989).

33 Shishkoff, N. & McGrath, M. AQ10 biofungicide combined with chemical fungicides or AddQ spray adjuvant for control of cucurbit powdery mildew in detached leaf culture. Plant Dis. 86, 915–918 (2002).

34 Romero, D., Rivera, M. E., Cazorla, F. M., De Vicente, A. & Perez-Garcia, A. Effect of mycoparasitic fungi on the development of *Sphaerotheca fusca* in melon leaves. Mycol. Res. 107, 64–71 (2003).

35 Siozios, S. et al. Transcriptional reprogramming of the mycoparasitic fungus *Ampelomyces quisqualis* during the powdery mildew host-induced germination. Phytopathology 105, 199–209 (2015). doi:10.1094/PHYTO-01-14-0013-R

36 Kiss, L. Natural occurrence of *Ampelomyces* intracellular mycoparasites in mycelia of powdery mildew fungi. New Phytol. 140, 709–714 (1998).

37 Szentiványi, O. et al. *Ampelomyces* mycoparasites from apple powdery mildew identified as a distinct group based on single-stranded conformation polymorphism analysis of the rDNA ITS region. Mycol. Res. 109, 429–438 (2005).

38 Liang, C. et al. Genetic diversity of *Ampelomyces* mycoparasites isolated from different powdery mildew species in China inferred from analyses of rDNA ITS sequences. Fungal Divers. 24, 225–240 (2007).

39 Park, M. J., Choi, Y. J., Hong, S. B. & Shin, H. D. Genetic variability and mycohost association of *Ampelomyces quisqualis* isolates inferred from phylogenetic analyses of ITS rDNA and actin gene sequences. Fungal Biol. 114, 235–247 (2010). doi:10.1016/j.funbio.2010.01.003

40 Kiss, L. et al. Temporal isolation explains host-related genetic differentiation in a group of widespread mycoparasitic fungi. Mol. Ecol. 20, 1492–1507 (2011). doi:10.1111/j.1365-294X.2011.05007.x

41 Pintye, A. et al. No indication of strict host associations in a widespread mycoparasite: grapevine powdery mildew (*Erysiphe necator*) is attacked by phylogenetically distant *Ampelomyces* strains in the field. Phytopathology 102, 707–716 (2012). doi:10.1094/PHYTO-10-11-0270

42 Angeli, D., Puopolo, G., Maurhofer, M., Gessler, C. & Pertot, I. Is the mycoparasitic activity of *Ampelomyces quisqualis* biocontrol strains related to phylogeny and hydrolytic enzyme production? Biol. Control 63, 348–358 (2012).

43 Pintye, A. et al. Host phenology and geography as drivers of differentiation in generalist fungal mycoparasites. PLoS One 10, e0120703 (2015). doi:10.1371/journal.pone.0120703

44 Brimner, T. A. & Boland, G. J. A review of the non-target effects of fungi used to biologically control plant diseases. Agric. Ecosyst. Environ. 100, 3–16 (2003). doi:10.1016/s0167-8809(03)00200-7

45 Kiss, L. How dangerous is the use of fungal biocontrol agents to nontarget organisms? New Phytol. 163, 453–455 (2004).

46 Sullivan, R. F. & White, J. F. *Phoma glomerata* as a mycoparasite of powdery mildew. Appl. Environ. Microbiol. 66, 425–427 (2000).

47 Aveskamp, M., De Gruyter, J., Woudenberg, J., Verkley, G. & Crous, P. W. Highlights of the *Didymellaceae*: a polyphasic approach to characterise *Phoma* and related pleosporalean genera. Stud. Mycol. 65, 1–60 (2010).

48 Szentivanyi, O. & Kiss, L. Overwintering of *Ampelomyces* mycoparasites on apple trees and other plants infected with powdery mildews. Plant Pathol. 52, 737–746 (2003).

49 Yarwood, C. E. An overwintering pycnidial stage of *Cicinnobolus*. Mycologia 31, 420–422 (1939).

50 Gorfer, M., Klaubauf, S., Bandian, D. & Strauss, J. *Cadophora finlandia* and *Phialocephala fortinii*: *Agrobacterium*-mediated transformation and functional GFP expression. Mycol. Res. 111, 850–855 (2007).

51 Kiss, L. Genetic diversity in *Ampelomyces* isolates, hyperparasites of powdery mildew fungi, inferred from RFLP analysis of the rDNA ITS region. Mycol. Res. 101, 1073–1080 (1997).

52 Angeli, D., Saharan, K., Segarra, G., Sicher, C. & Pertot, I. Production of *Ampelomyces quisqualis* conidia in submerged fermentation and improvements in the formulation for increased shelf-life. Crop Protect. 97, 135–144 (2017). doi:10.1016/j.cropro.2016.11.012

53 Marboutie, G., Combe, F., Boyer, E. & Berne, A. Travaux sur l’*Ampelomyces quisqualis* (1993-1994). IOBC/WPRS Bulletin 18, 75–78 (1995).

54 Idnurm, A. et al. A silver bullet in a golden age of functional genomics: the impact of *Agrobacterium*-mediated transformation of fungi. Fungal. Biol. Biotechnol. 4, 6 (2017).

55 Li, M. et al. Transformation of *Coniothyrium minitans*, a parasite of *Sclerotinia sclerotiorum*, with *Agrobacterium tumefaciens*. FEMS Microbiol. Lett. 243, 323–329 (2005).

56 Katoh, K. & Standley, D. M. MAFFT multiple sequence alignment software version 7: improvements in performance and usability. Mol. Biol. Evol. 30, 772–780 (2013). doi:10.1093/molbev/mst010

57 Tamura, K., Stecher, G., Peterson, D., Filipski, A. & Kumar, S. MEGA6: Molecular Evolutionary Genetics Analysis version 6.0. Mol. Biol. Evol. 30, 2725–2729 (2013). doi:10.1093/molbev/mst197

58 Silvestro, D. & Michalak, I. raxmlGUI: a graphical front-end for RAxML. Org. Divers. Evol. 12, 335–337 (2012).

59 Stamatakis, A. RAxML version 8: a tool for phylogenetic analysis and post-analysis of large phylogenies. Bioinformatics 30, 1312–1313 (2014).

60 Stöver, B. C. & Müller, K. F. TreeGraph 2: Combining and visualizing evidence from different phylogenetic analyses. BMC Bioinformatics 11, 7 (2010). doi:10.1186/1471-2105-11-7

61 Hanif, M., Pardo, A., Gorfer, M. & Raudaskoski, M. T-DNA transfer and integration in the ectomycorrhizal fungus *Suillus bovinus* using hygromycin B as a selectable marker. Curr. Genet. 41, 183–188 (2002).

62 Lazo, G. R., Stein, P. A. & Ludwig, R. A. A DNA transformation-competent *Arabidopsis* genomic library in *Agrobacterium*. Nat. Biotechnol. 9, 963–967 (1991).

63 Carroll, A. M., Sweigard, J. A. & Valent, B. Improved vectors for selecting resistance to hygromycin. Fungal. Genet. Rep. 41, 22 (1994).

64 Cubero, O. F., Crespo, A., Fatehi, J. & Bridge, P. D. DNA extraction and PCR amplification method suitable for fresh, herbarium-stored, lichenized, and other fungi. Plant Syst. Evol. 216, 243–249 (1999).

65 Kumar, S., Stecher, G. & Tamura, K. MEGA7: Molecular Evolutionary Genetics Analysis version 7.0 for bigger datasets. Mol. Biol. Evol. 33, 1870–1874 (2016).

66 Livak, K. J. & Schmittgen, T. D. Analysis of relative gene expression data using real-time quantitative PCR and the 2^-AACT^ method. Methods 25, 402–408 (2001).

67 Bubner, B. & Baldwin, I. T. Use of real-time PCR for determining copy number and zygosity in transgenic plants. Plant Cell Rep. 23, 263–271 (2004).

68 Solomon, P. S., Ipcho, S. V., Hane, J. K., Tan, K.-C. & Oliver, R. P. A quantitative PCR approach to determine gene copy number. Fungal. Genet. Rep 55, 5–8 (2008).

69 Gorfer, M. et al. Community profiling and gene expression of fungal assimilatory nitrate reductases in agricultural soil. ISME J. 5, 1771–1783 (2011).

70 Schneider, C. A., Rasband, W. S. & Eliceiri, K. W. NIH Image to ImageJ: 25 years of image analysis. Nat. Methods 9, 671–675 (2012).

71 Vági, P. et al. Simultaneous specific in planta visualization of root-colonizing fungi using fluorescence in situ hybridization (FISH). Mycorrhiza 24, 259–266 (2014).

